# Hypoxia-induced miR-92a regulates p53 signalling pathway and apoptosis by targeting calcium-sensing receptor in Genetically Improved Farmed Tilapia (*Oreochromis niloticus*)

**DOI:** 10.1101/2020.08.27.269670

**Authors:** Jun Qiang, Jie He, Yi-Fan Tao, Jin-Wen Bao, Jun-Hao Zhu, Pao Xu

## Abstract

miR-92a miRNAs are immune molecules that regulate apoptosis (programmed cell death) during the immune response. Apoptosis helps to maintain the dynamic balance in tissues of fish under hypoxia stress. The aim of this study was to explore the role and potential mechanisms of miR-92a in the liver of tilapia under hypoxia stress. We first confirmed that *CaSR* (encoding a calcium-sensing receptor) is a target gene of miR-92a in genetically improved farmed tilapia (GIFT) using luciferase reporter gene assays. In GIFT under hypoxia stress, miR-92a was up-regulated and *CaSR* was down-regulated in a time-dependent manner. Knocked-down *CaSR* expression led to inhibited expression of *p53*, *TP53INP1* and *caspase-3/8*, reduced the proportion of apoptotic hepatocytes, and decreased the activity of calcium ions induced by hypoxia in hepatocytes. GIFT injected in the tail vein with an miR-92a agomir showed up-regulation of miR-92a and down-regulation of *CaSR*, *p53*, *TP53INP1*, and *caspase-3/8* genes in the liver, resulting in lower serum alanine aminotransferase and aspartate aminotransferase activities under hypoxia stress. These findings suggest that stimulation of miR-92a interferes with hypoxia-induced apoptosis in hepatocytes of GIFT by targeting *CaSR*, thereby alleviating liver damage. These results provide new insights into the adaptation mechanisms of GIFT to hypoxia stress.

## Background

miRNAs are a small class of non-coding RNAs that regulate the expression of one or more target genes by binding to their 3′-untranslated region (UTR). In recent years, the potential application of members of the miR-92 family as immune molecules in aquaculture has received extensive attention (Zhang et al., 2014; Chen et al., 2016; Yang et al., 2013; Qiang et al., 2017). Previous studies have shown that the miR-92 family is closely related to the immune response of the sea cucumber *Apostichopus japonicus* to vibriosis caused by *Vibrio splendidus* (Zhang et al., 2014). That study showed that miR-92 as regulate the host–pathogen interaction in sea cucumber by binding to two candidate genes, one encoding polyepidermal growth factor-like domain 6 and the other encoding SMAD-specific E3 ubiquitin protein ligase. In the oyster *Crassostrea gigas*, miR-92d regulates the expression of tumour necrosis factor (TNF) by targeting the coding region of *CgLITAF3*, which encodes lipopolysaccharide-induced TNF-a factor 3, thereby triggering an inflammatory response to invading bacteria (Chen et al., 2016). In amphioxus *Branchiostoma belcheri*, miR-92d regulates the immune response to bacterial infection by binding to the gene encoding complement C3 (Yang et al., 2013). In genetically improved farmed tilapia (GIFT, *Oreochromis niloticus*), miR-92d-3p is involved in mediating the expression of complement C3, and inhibition of miR-92d-3p promotes complement C3 expression and enhances the inflammatory response to *Streptococcus iniae* infection (Qiang et al., 2017).

In a previous study, we successfully constructed miRNA expression libraries from uninfected GIFT and those infected with *S. iniae* (Qiang et al., 2017). Gene enrichment pathway analyses indicated that one of the differentially expressed miRNAs, miR-92a, may be involved in cell signal transduction or immune regulation processes in GIFT (Qiang et al., 2017). miR-92a family play important roles in the regulation of cell tumour proliferation, apoptosis, invasion, and metastasis (Niu et al., 2012; Tsuchida et al., 2011; Ohyagi-Hara et al., 2013). A previous study showed that miR-92a is highly expressed in the tumour tissue of patients with glioblastoma, and that its expression level is negatively correlated with the level of Bcl-2 interacting mediator of cell death (Bim) and positively correlated with tumour malignancy (Niu et al., 2012). miR-92a directly targets the *Bim* gene. Application of miR-92a inhibitors can significantly increase tumour cell apoptosis, suggesting that miR-92a has a role in promoting tumorigenesis. Tsuchida et al (2011) found that miR-92a is highly expressed in colon adenoma and colon cancer cells, and it targets the *Bim* gene to inhibit apoptosis of cancer cells. Application of an miR-92a antagomir was shown to affect the rate of apoptosis of colon cancer cell lines.

miRNAs are highly conserved among different species and are genetically stable. Some important miRNA signalling pathways may have the same or similar roles in fish as in mammals (Qiang et al., 2017). *CaSR* (encoding a calcium-sensing receptor) may be a potential target gene of miR-92a (free energy, −26.5 Kcal/mol) in GIFT. In mammals, CaSR is a sensitive receptor for Ca^2+^, and its expression is regulated by the concentration of extracellular Ca^2+^. As a multifunctional regulator, CaSR participates in G protein signal transduction and GPCRs kinase (GPK)-induced desensitisation by altering the intracellular Ca^2+^ concentration, thereby mediating processes such as cell growth, differentiation, and ion channel opening (Díaz-Soto et al., 2016; Vezzoli et al., 2019). To expand on the results of those studies and our previous studies, we aimed to confirm the relationship between miR-92a and its target *CaSR*, and determine how this relationship regulates cell stress responses in GIFT. We also aimed to clarify its regulatory pathway and activation mechanism. Addressing these questions will shed light on the molecular mechanisms of stress regulation in fish.

Changes in dissolved oxygen (DO) can affect fish survival. Generally, fish grow and develop normally when the DO level is higher than 4.0 mg/L. Low DO levels (<2.0 mg/L) cause a floating phenomenon in farmed fish. If the DO level drops below 1.0 mg/L (hypoxic conditions), most fish will severely float their heads and eventually suffocate to death (Perry et al., 2011). There are many causes of fish death under hypoxia stress, including an imbalance of apoptosis (Sun et al., 2020). Apoptosis refers to the orderly death of cells controlled by genes to maintain the stability of the internal environment. Apoptosis and the immune response involve proteins including p53, p53-inducible nuclear protein 2 (TP53INP2), caspase-3, and caspase-8. Among them, p53 is considered to be a key factor in the regulation of apoptosis, because it directly or indirectly induces multiple regulatory genes to interfere with apoptosis (Gong et al., 2015). As a downstream gene of p53, *TP53INP* encodes a protein that coactivates transcription factors such as p53 in the nucleus to regulate apoptosis and the expression of cell cycle-related genes (Tomasini et al., 2005). Apoptosis-associated caspases are widely expressed in various fish tissues, with relatively high expression levels in immune-related tissues (Denko et al., 2003). The expression levels of *caspase-3* and *caspase-8* were found to be low in healthy sea bass (*Dicentrarchus labrax*) but increased after infection with *Photobacterium damselae ssp.* (Reis et al., 2007; 2010). Similarly, in large yellow croaker (*Pseudosciaena crocea*), the expression levels of *caspase-3* and *caspase-9* in the kidney and spleen were found to be significantly increased after stimulation with poly I:C and bacteria, and the apoptotic pathway was activated (Mu et al., 2010; Li et al., 2011).

Tilapia is one of the most important freshwater farmed fish in China. High temperatures or deterioration of water quality can cause a sharp drop in DO levels in aquaculture ponds, leading to fish death (Yuan et al., 2019). Therefore, the main objectives of this study were as follows: 1) to verify that *CaSR* is a potential target gene of miR-92a; 2) to determine the effect of *CaSR* knockdown on the Ca^2+^ concentration and apoptosis pathway in GIFT hepatocytes; and 3) to explore how miR-92a expression is activated in GIFT under hypoxia stress. The results of this study provide new information about stress regulation and adaptation mechanisms in fish.

## 2. Materials and methods

### Ethics statement

The study protocols and design were approved by the Ethics Committee at the Freshwater Fisheries Research Center of the Chinese Academy of Fishery Sciences (Wuxi, China). The GIFT were maintained in well-aerated water and treated with 200 mg/L tricaine methane sulfonate (Sigma, St Louis, MO, USA) for rapid deep anaesthesia. The samples were extracted based on the Guide for the Care and Use of Laboratory Animals in China.

### Identification of binding sites of miR-92a-CaSR 3′UTR

The full-length *CaSR* 3′ UTR sequence (XM_025910710.1) of GIFT was synthesised and then inserted into the pGL3-control vector, yielding 3′ UTR wild-type. In the seed sequence region where miRNA binds to its target gene, the six complementary bases (AUUGUA) of CaSR 3′-UTR were replaced with GACAAG to construct the 3′UTR mutant. The procedures for HEK293T cell culture, vector transfection, and luciferase activity determination were as described previously (Qiang et al., 2018). We used Renilla luciferase activity to standardize the luciferase activity of each transfected cell. As the miRNA negative control (NC), we selected the miRNA of *Caenorhabditis elegans*, which has minimal homology with all miRNAs in the miRBase database. There were four treatment groups in this experiment: CaSR 3′ UTR-wild type + miRNA mimic; CaSR 3′ UTR-Mutant + miRNA mimic; miRNA NC (Ctrl mimic) + CaSR 3′ UTR-wild type; Ctrl mimic + CaSR 3′ UTR-Mutant. Each treatment had eight replicates. Luciferase activity was detected to determine the binding efficiency of miRNA to its target gene.

### Analysis of the regulatory relationship between miRNA and its target genes

One hundred and eighty healthy GIFT (average fish weight: 18.4 g ± 0.8 g) were randomly assigned to nine experimental groups (20 fish/group). There were three treatment groups: miR-92a agomir group; miR-92a negative agomir group; PBS group (three replicates of 20 fish/treatment). The chemically synthesised miRNA agomir and miRNA negative agomir (with the six mismatch mutation in its complementary bases) were designed and synthesised by the Ruibo Biological Technology Co., Ltd. (Guangzhou, China). Every 6 days, miRNA agomir was administered by tail vein injection at a dose of 25 mg/kg body weight. The same dose of NC or PBS was administered to fish in the NC or PBS groups, respectively, in the same way. At 0 h, 12 h, 24 h, and 48 h post-injection, three fish were randomly selected from each tank and deeply anesthetized before dissecting liver tissues. The liver samples were frozen in liquid nitrogen within 1 h, and then stored at −80 °C until analyses of gene expression and protein levels.

### Regulatory response of GIFT hepatocytes under hypoxia stress after knocking down expression of *CaSR*

We constructed an RNA interference (RNAi) expression vector to knock down the expression of *CaSR* in GIFT. First, according to the GIFT *CaSR* gene sequence, we designed an RNAi (5′-GAGAGCACAGATGACTGGTGATATT-3′) sequence with no homology to other coding sequences of tilapia. The construction of the positive interference plasmid (CaSR knockdown), the negative control interference plasmid (NC), and the control plasmid (Con) were as described in Qiang et al. (2020). Hepatocytes were cultured, isolated, and purified as described by Chen et al (2011). This experiment had four experimental groups of cells: hypoxia treatment (CaSR knockdown group, NC group, and Con group), and normal group (NG). Each treatment had eight replicates. The primary cultured hepatocytes were grown to about 80%–90%, and then transfected using Lipofectamine 2000. The three hypoxia-treated cell groups were placed in an YQX-II anoxic incubator (JTONE, Hangzhou, China) (N_2_ 93%, O_2_ 2%, CO_2_ 5%) for 24 h to impose hypoxia stress. The cell lysate was treated for 30 min, and then centrifuged (12000 *g*, 20 min, 4 °C). The supernatant was collected for further analyses. The hepatocytes in the NG were cultured normally without transfection and hypoxia stress.

### miR-92a-mediated stress response in the liver of GIFT after hypoxia exposure

The experimental design was based on that of Qiang et al. (2018). In total, 360 GIFT juveniles (average size, 26.8 g ± 0.9 g) were randomly assigned to nine 400-L tanks (40 fish/tank). The synthesis, dissolution, and injection of the miR-92a agomir and miRNA NC were as described in section 2.2. The control group was similarly injected with PBS. The experimental GIFT at 12 h post-injection were subjected to low-DO (1.0 mg/L) stress for 96 h. The DO level in water was determined using an LDO101 probe (Hach, Loveland, CO, USA). Nitrogen and air charges were used to regulate and maintain the DO level in water. Three fish were randomly selected from each group at each sampling time point, and blood was drawn and liver tissues were dissected quickly. The liver tissues were stored at −80 °C until analyses of mRNA levels. Blood samples were left at 4 °C for 2 h and then centrifuged (4 °C, 3500 *g*, 10 min) to collect the serum. The serum was kept at −80 °C until analysis.

### Measurement indices

#### mRNA expression

Total RNA extraction, reverse transcription (RT), and quantitative real-time PCR (qRT-PCR) were as described elsewhere (Qiang et al., 2020). qRT-PCR (see Table 1 for primers) analyses were conducted using the ABI QuantStudio 5 Real-Time PCR System (Foster City, CA, USA), with *18S rRNA* as the endogenous control (reference gene).

**Table 1.**
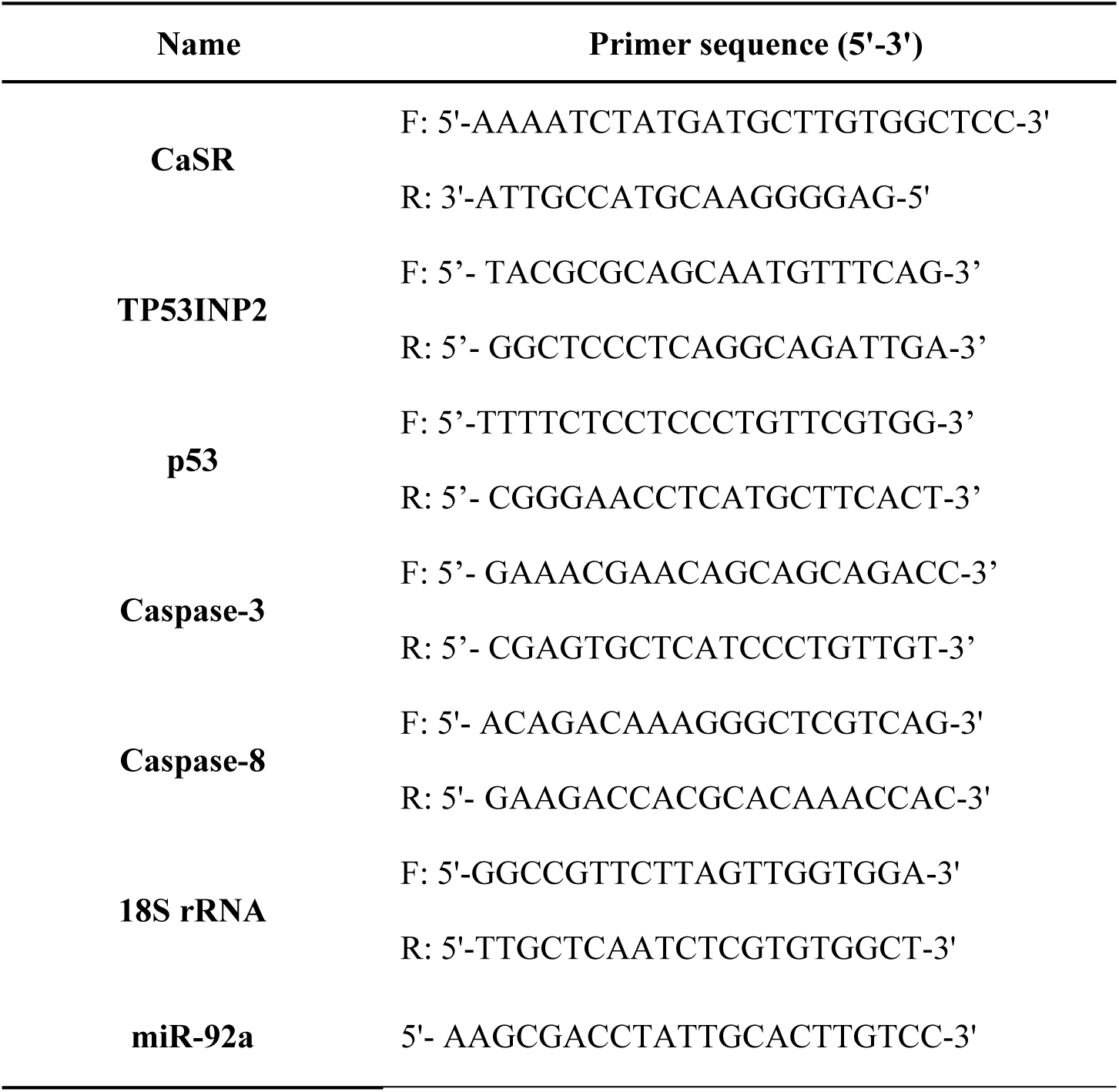
Sequences of primers used for qRT-PCR

#### miRNA expression

The RT reaction and qRT-PCR of the miRNA were as described elsewhere (Qiang et al., 2020). The primers for miR-92a are shown in Table 1. The reference gene was *U6*.

#### Western blot analysis

Each liver tissue sample (0.1 g) was ground in liquid nitrogen with a mortar and pestle, then 1 mL ristocetin-induced platelet agglutination buffer (containing 1% v/v 10 mg/ml phenylmethanesulfonyl fluoride) was added and the mixture was homogenized (15,000 g, 1 min, 4 °C) using a Polytron (PT2500E, Kinematica, Lucerne, Switzerland). The procedures for SDS-PAGE preparation, protein sample electrophoresis, membrane transfer, blocking, and antibody incubation were as described by Qiang et al. (2018). Colour was developed using Immobilon Western HRP substrate (Millipore, Billerica, MA, USA). Glyceraldehyde-3-phosphate dehydrogenase (GAPDH) was the reference protein.

#### Identification of apoptotic cells and determination of Ca^2+^ concentration

The culture, collection, and apoptotic cell detection of hepatocytes were as described by Qiang et al. (2020). Hepatocytes were collected and resuspended in pre-cooled PBS, and then the Ca^2+^-fluorescent probe Fluo-3AM was added to a final concentration of 5 μg/mL. The mixture was incubated in the dark for 40 min at room temperature, and then passed through a cell sieve (40 μm mesh size). The Ca^2+^ concentration was detected using the Incyte module of a FACSCalibur flow cytometer with excitation at 506 nm and emission at 526 nm.

#### Measurement of activities of alanine aminotransferase (ALT) and aspartate aminotransferase (AST) in serum

The activities ALT and AST in serum were determined using kits obtained from the Jiancheng Bio-Engineering Institute (Nanjing, China), according to the manufacturer’s instructions. Absorbance at the wavelengths specified in the kit manuals was determined using a BioTek Eon Microplate Spectrophotometer (BioTek, Winooski, VT, USA).

### Data processing and statistical analysis

The relative expression levels of mRNA and miRNA were calculated using the 2-ΔΔCT method. Data from all experiments were statistically analysed using SPSS 21.0 software. Shapiro–Wilk’s test was used to test whether the data conformed to a normal distribution (α=0.1). Levene’s test was used to determine the homogeneity of variance (α=0.1). The experimental data shown in figures and tables are mean ± standard deviation. When the experimental data met the tests for normality and homogeneity of variance, we performed analysis of variance and selected appropriate statistical methods for comparisons among treatment groups.

## Results

### Verification of binding site and regulatory relationship between miR-92a and its target gene

We generated an miR-92a mimic to target the 3′ end of *CaSR*, and then tested its efficacy by conducting dual luciferase reporter assays using HEK293T cells. Luciferase activity in the *CaSR* 3′ UTR wild type + control mimic group (Ctrl group, no mimic) was 1.12 ± 0.03, while that in the UTR wild type + miR-92a mimic group was decreased by 48.47% (Figure 1A) (statistical significance; t = 2.357, *P* = 0.023). To confirm that binding between the *CaSR* 3′ UTR and the miR-92a sequence was essential for this activity, we constructed a mutated *CaSR* 3′ UTR sequence (3′ UTR Mut). The luciferase activity levels in the 3′ UTR Mut + Ctrl mimic group or the miR-92a mimic group were 0.96-times and 0.87-times that in the control group, respectively; these differences were not significant (*P* > 0.05). These results confirmed that GIFT miR-92a binds to the 3′ UTR of *CaSR* mRNA. Although the 5′ end of the miR-92a sequence showed only partial complementarity in a 2–8 bp region, a compensation site at the 3′ end enhanced its binding stability to the 3′-UTR of the target *CaSR* mRNA (Figure 1B).

**Figure 1.**
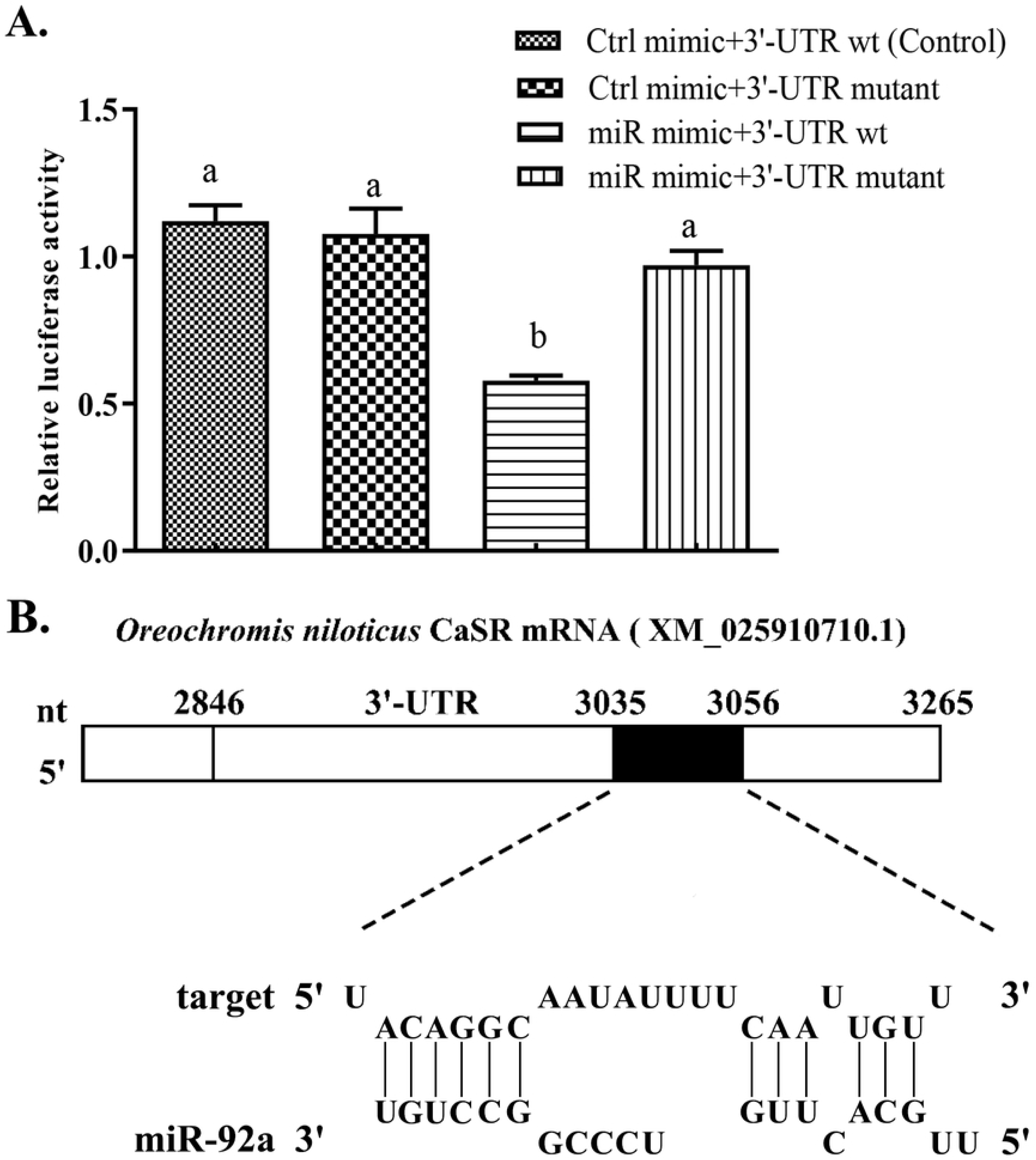
Verification of binding between miRNA and its potential target gene. (A) Validation of miRNA binding to 3′ UTR of potential target gene using dual luciferase reporter system. HEK-293T cells in 12-well plates were co-transformed with pGL-CaSR 3′ UTR (3′ UTR-wild type) or pGL-CaSR (3′ UTR-Mutant) and miRNA mimic (miR mimic) or miRNA negative control (Ctrl mimic) using Lipofectamine 2000 transfection reagent. Luciferase activity was determined based on fluorescence. Different lowercase letters indicate significant differences among experimental groups (Duncan’s multiple comparison; *P* < 0.05). (B) Analysis of binding site between miR-92a and *Oreochromis niloticus CaSR* (XM_025910710.1): 5′-end of miR-92a can pair with 3035 bp–3056 bp position at the *CaSR* 3′-UTR. A compensation binding site at the 3′-end of miR-92a enhances its binding stability with 3′-UTR of *CaSR*.

To analyse the regulatory relationship between miR-92a and its target gene, we injected the miR-92a agomir. negative control (NC) or PBS (control) into the tail vein of GIFT. The levels of hepatic *CaSR* mRNA and protein were significantly lower in GIFT treated with the miRNA agomir than in the control group at 12, 24, and 48 h after injection (Figure 2A and Figure S1, *P* < 0.05); however, the levels of miR-92a were significantly increased (Figure 2B). There was no significant difference in the levels of *CaSR* mRNA and miR-92a between the control group and the NC group at each time point. These results showed that up-regulation of miR-92a expression led to down-regulation of *CaSR*. *CaSR knockdown regulated Ca^2+^ concentration and expression of genes related to the cell signalling pathway in GIFT hepatocytes*

**Figure 2.**
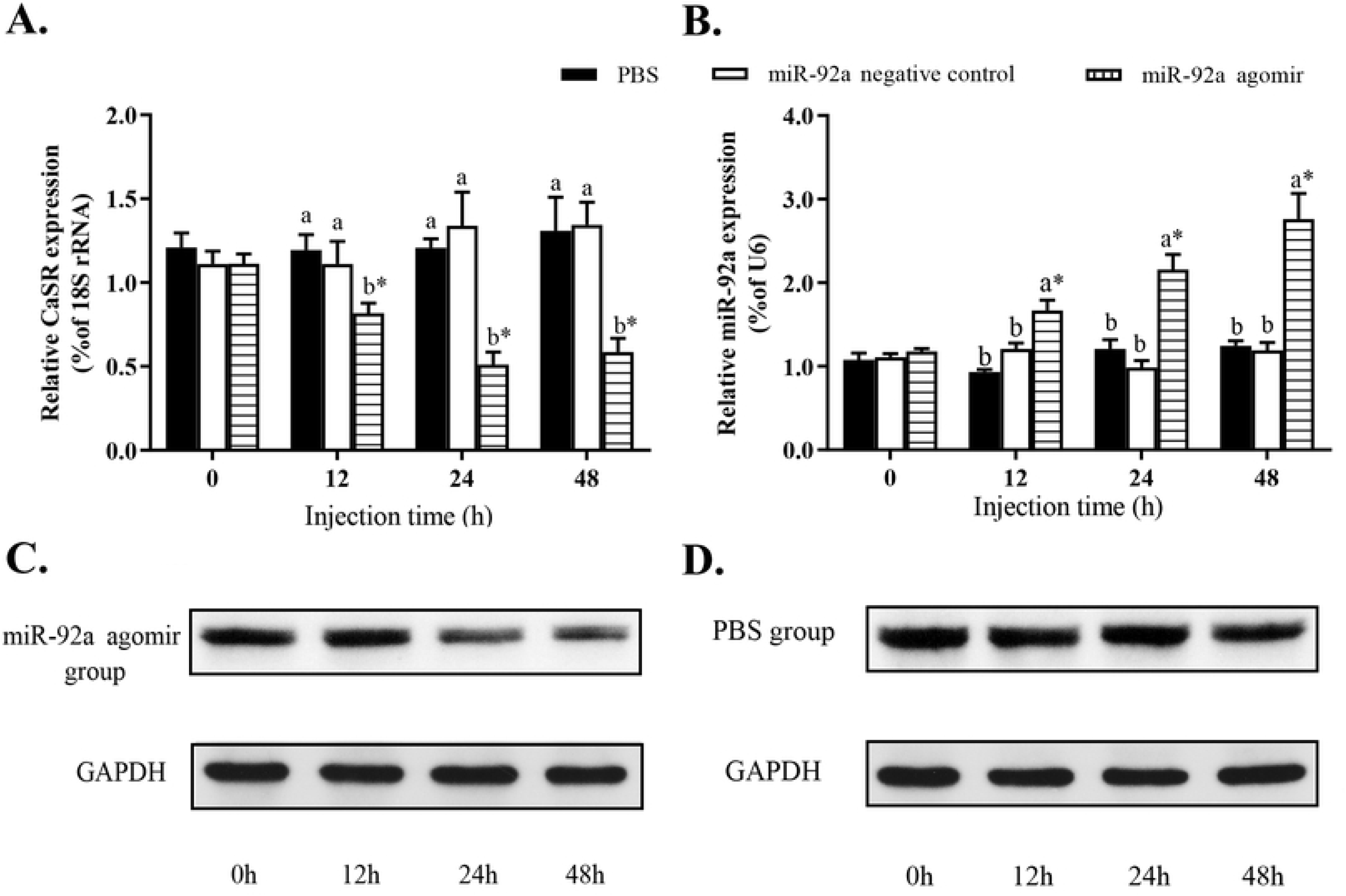
Analysis of regulatory relationship between *CaSR* (A) and miR-92a (B) *in vivo*. Juvenile GIFT weighing about 18.4 g ± 0.8 g were injected in tail vein with miR-92a negative control, miR-92a agomir (dose, 25 mg/kg body weight), or PBS (control). (A, C and D) qRT-PCR (upper panel) and western blot (lower panel) analyses of CaSR expression in GIFT after injection with miR-92a agomir. Bands of interest were cropped from the same membrane. (B) qRT-PCR analysis of miR-92a expression in GIFT after injection with miR-92a agomir. Based on relative levels in control group at 0 h, relative expression levels of mRNA and miRNA in each experimental group were determined by 2^−ΔΔCT^ method. * indicates significant differences between values before and after injection (paired-samples t test; *P* < 0.05). Different lowercase letters show significant differences among different treatments at each sampling point (Duncan’s multiple range test; *P* < 0.05).

Next, we determined the effects of CaSR knockdown on the responses of GIFT hepatocytes under hypoxia stress. This experiment consisted of four groups of GIFT hepatocytes: a normal group (NG, no transfection, no hypoxia); a control + hypoxia stress group (Con + hypoxia); a CaSR-negative control transfection + hypoxia stress group (NC + hypoxia); and a CaSR knockdown + hypoxia stress group (Knockdown + hypoxia).

As shown in Figure 3A, compared with the *CaSR* transcript levels in the Con + hypoxia group and the NC + hypoxia group, that in the Knockdown + hypoxia group was decreased by 60% (*P* < 0.05). The transcript levels of *TP53INP2* (Figure 3B), *p53* (Figure 3C), *caspase-3* (Figure 3D), and *caspase-8* (Figure 3E) were significantly lower in the Knockdown+ hypoxia group than in the Con + hypoxia and NC + hypoxia groups (Figure 3B). The percentage of hepatocytes in the early and late stage of apoptosis was 14.62% in the Con + hypoxia group and 14.28% in the NC + hypoxia group (Q4 +Q2 in Figure 4A), significantly higher than in the NG and Knockdown + hypoxia group (Figure 4A). The Ca^2+^ concentration was significantly lower in the Knockdown + hypoxia group than in the Con + hypoxia and NC + hypoxia groups (Figure 4B), but not significantly different between the NG and the knockdown+ hypoxia group (*P*>0.05).

**Figure 3.**
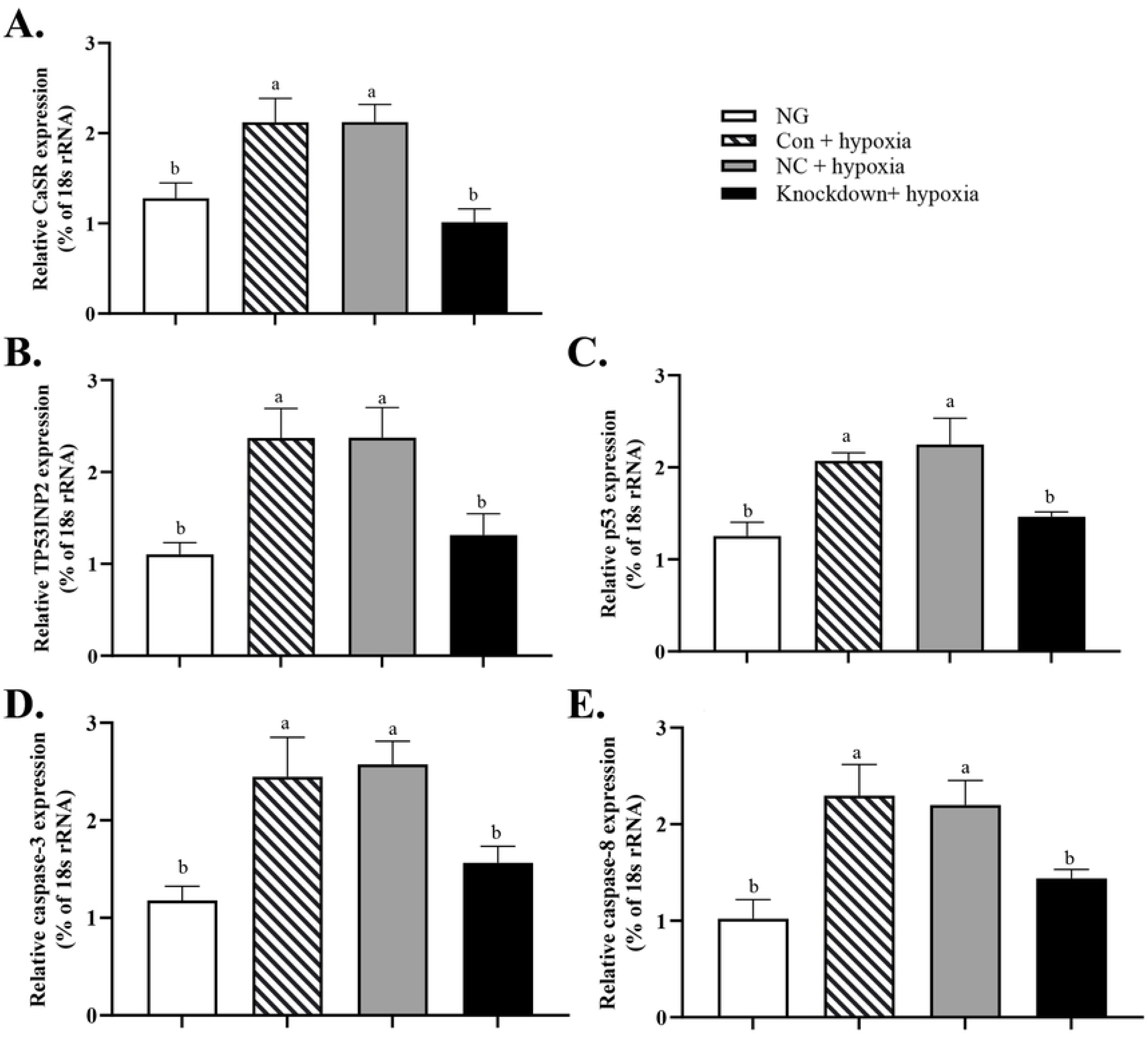
Effect of *CaSR* knockdown on transcript levels of *CaSR*, *p53*, *p53-inducible nuclear protein 2*, *caspase-3*, and *caspase-8* in GIFT hepatocytes under hypoxia stress. (A) Effect of *CaSR* knockdown on transcript levels of CaSR in GIFT hepatocytes under hypoxia stress. Hepatocytes were transfected with *CaSR* knockdown, NC, or Con vector using Lipofectamine 2000 and subjected to 24 h of hypoxia stress in an incubator (knockdown+ hypoxia; NC+ hypoxia, and Con+ hypoxia, respectively). Normal group (NG): hepatocytes without transfection and without hypoxia. (B, C, D and E) Effect of *CaSR* knockdown on transcript levels of *p53*, *p53-inducible nuclear protein 2*, *caspase-3* and *caspase-8* in GIFT hepatocytes under hypoxia stress. Based on relative levels of respective mRNAs in NG, relative mRNA levels in each group were determined by 2-ΔΔCT method. Different lowercase letters indicate significant differences among different treatments (Duncan’s multiple range test, *P* < 0.05).

**Figure 4.**
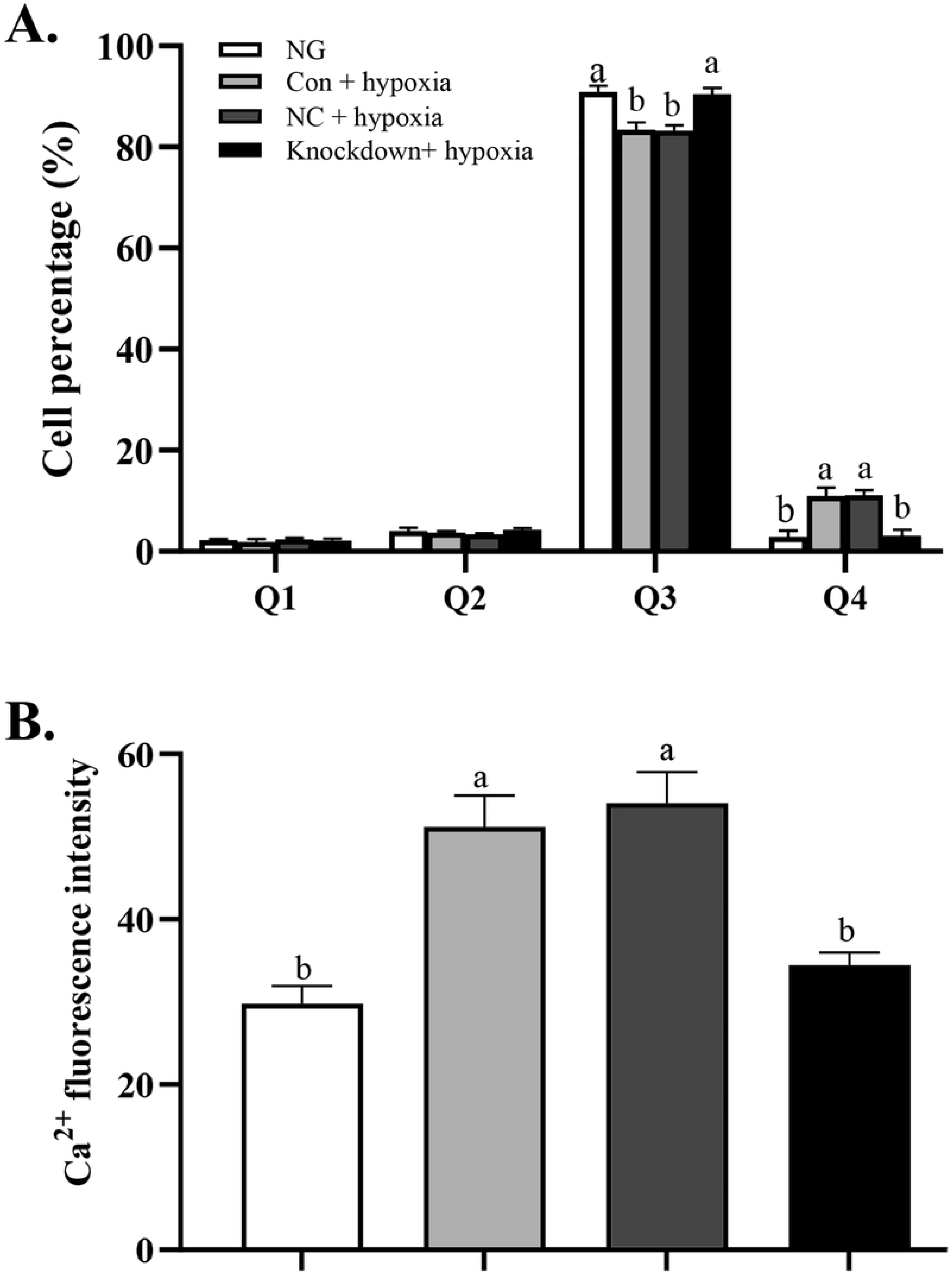
Effect of *CaSR* knockdown on apoptosis and Ca^2+^ concentration in hepatocytes of GIFT under hypoxia stress. (A) Proportion of apoptotic cells in NG, knockdown+ hypoxia, NC+ hypoxia and Con+ hypoxia groups. Figure shows percentage of dead cells (Q1), late apoptotic cells (Q2), viable cells (Q3), and early apoptotic cells (Q4). (B) Ca^2+^ fluorescence intensity in NG, knockdown+ hypoxia, NC+ hypoxia and Con+ hypoxia groups. Different lowercase letters indicate significant differences among different treatments (Duncan’s multiple range test, *P* < 0.05).

### miR-92a mediated participation of target gene in hepatic stress regulation in GIFT under hypoxia stress

To further analyse the role of miRNA-mediated *CaSR* expression in the GIFT liver in response to hypoxia stress, juveniles were injected with miR-92a agomir via the tail vein. At 12 h after miRNA agomir injection, the GIFT were subjected to hypoxia stress. The transcript level of *CaSR* was significantly lower in the agomir-treated group than in the control group at 12, 24, 48, 72, and 108 h of hypoxia stress (Figure 5B). At each time point, the level of miR-92a was significantly higher in the agomir group than in the control and the NC groups (Figure 5A).

**Figure 5.**
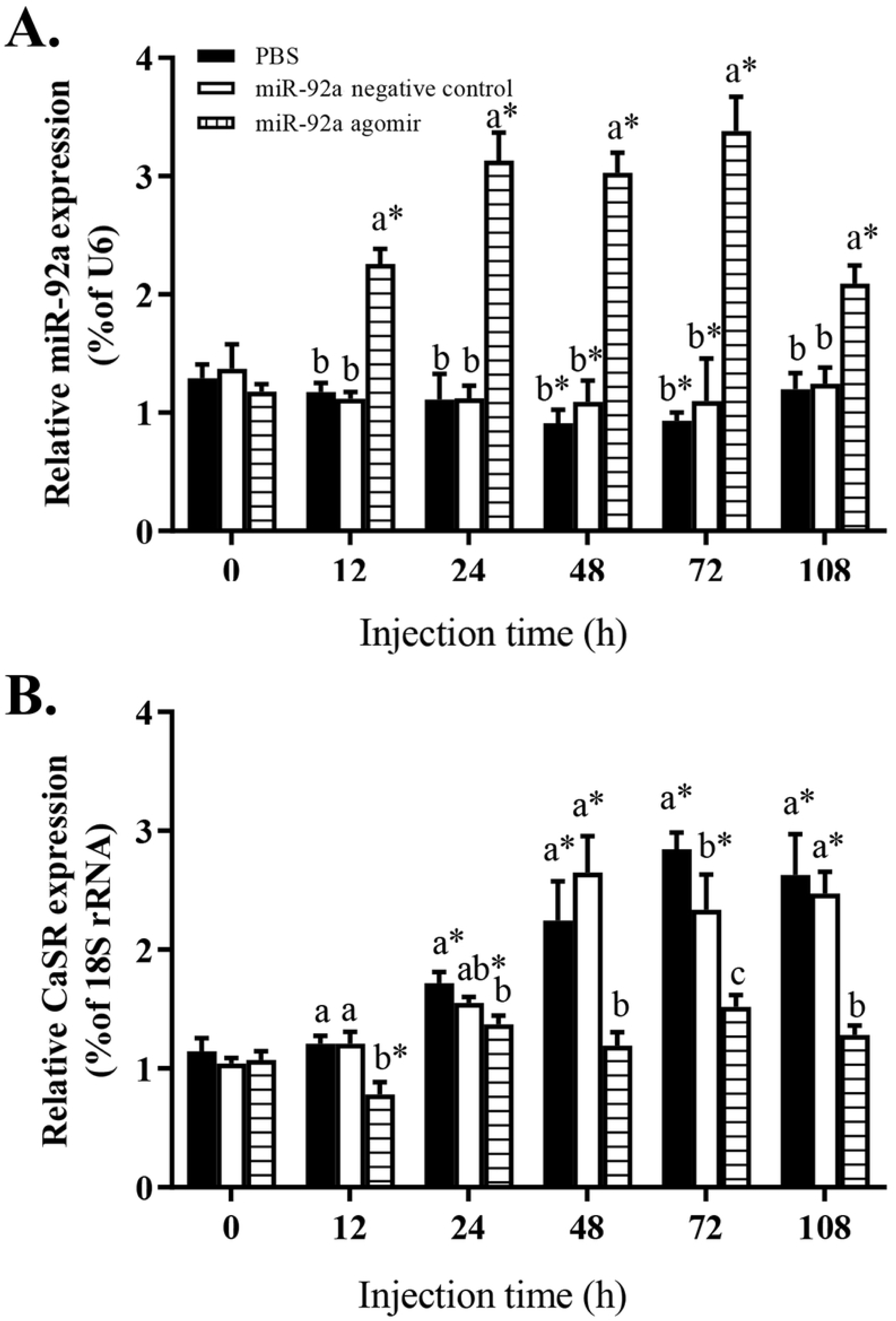
Expression levels of miR-92a (A) and *CaSR* (B) in liver of GIFT treated with miRNA agomir to promote miR-92a expression. Juveniles weighing about 26.8 g ± 0.9 g were injected in tail vein with miR-92a negative control, or miR-92a agomir (dose, 25 mg/kg body weight) or PBS (control) and response was monitored for 108 h. Based on relative expression levels in control group at 0 h, relative expression levels of mRNA and miRNA in each experimental group were determined by 2^−ΔΔCT^ method. * indicates significant differences between before and after injection (paired-samples t test; *P* < 0.05). Different lowercase letters indicate significant differences among different treatments at each sampling point (Duncan’s multiple comparison; *P* < 0.05).

After 24 h of hypoxia stress, the serum alanine aminotransferase (ALT) and aspartate (AST) aminotransferase activities were significantly lower in the agomir group than in the control and the NC groups (Figure 6). The serum AST and ALT activities in all treatment groups tended to increase under hypoxia stress. Serum ALT activity in the agomir group was significantly higher at 24 and 48 h than at 0 h (pre-stress). The transcript levels of *p53, TP53INP2, caspase-3*, and *caspase-8* in the liver tended to increase under hypoxia stress in the control group and the NC group (Figure 7). The transcript levels of *caspase-3* were significantly lower in the agomir group than in the control group and the NC group at 24, 48, and 96 h of hypoxia stress. The transcript levels of *p53* and *caspase-8* were lower in the agomir group than in the control group at 96 h of hypoxia stress. Additionally, the transcript level of *TP53INP2* was significantly lower in the agomir group than in the control group at 48 and 96 h of hypoxia stress.

**Figure 6.**
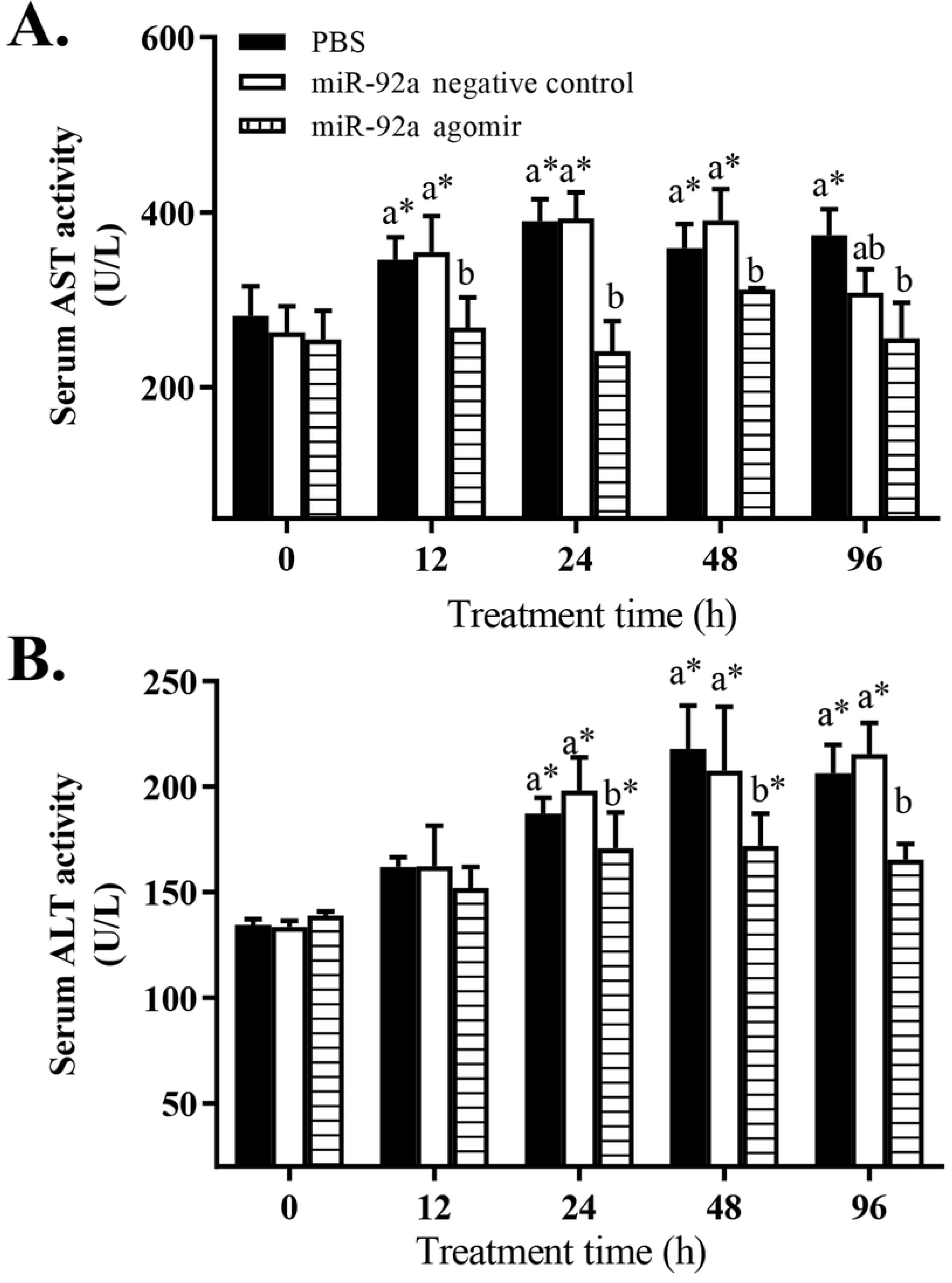
Effect of increased miR-92a expression on serum alanine aminotransferase (A) and aspartate aminotransferase (B) activities in GIFT under hypoxia stress. At 12 h after miR-92a agomir injection, juveniles were subjected to hypoxia stress for 96 h. Juveniles injected with PBS served as control. * indicates significant difference between before and after injection (paired-samples t test; *P* < 0.05). Different lowercase letters show significant differences among different treatments at each sampling point (Duncan’s multiple comparison; *P* < 0.05).

**Figure 7.**
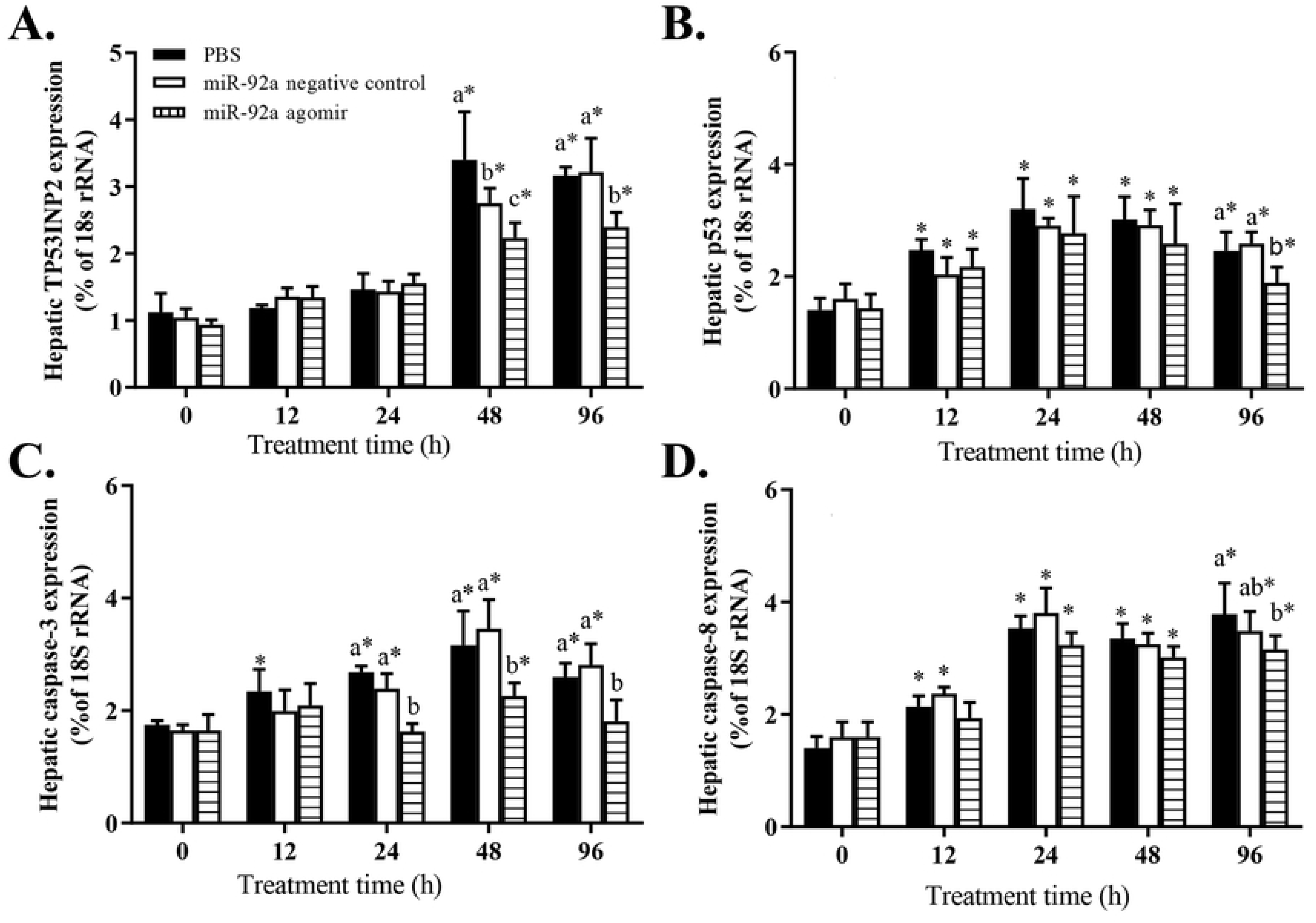
Effect of promoting miR-92a expression on transcript levels *p53-inducible nuclear protein 2* (A), *p53* (B), *caspase-3* (C), and *caspase-8* (D) in GIFT under hypoxia stress. At 12 h after injection with miR-92a agomir or PBS (control), juveniles were subjected to hypoxia stress for 96 h. Based on relative expression level in control group at 0 h, relative expression levels of mRNAs in each experimental group were determined by 2^−ΔΔCT^ method. * indicates significant differences between before and after injection (paired-samples t test; *P* < 0.05). Different lowercase letters indicate significant differences among different treatments at each sampling point (Duncan’s multiple comparison; *P* < 0.05).

## Discussion

In fish, CaSR is involved in cellular pro-inflammatory responses and apoptosis signalling (Klein et al., 2016; Hernández-Bedolla et al., 2016). Calcium ions are second messengers of signal transduction and can trigger a variety of cellular and physiological responses, such as muscle contraction, exocytosis, neurotransmitter release, cell proliferation, and apoptosis (Klein et al., 2016; Hendy et al., 2016). In this study, the Ca^2+^ concentration in GIFT hepatocytes increased sharply under hypoxia stress, which may explain the increased proportions of cells in the early and late stages of apoptosis. Previous studies have shown that an increase in the Ca^2+^ concentration induces expression of pro-inflammatory cytokines, thereby activating caspase-9 in the mitochondrial pathway or caspase-8 in the death receptor pathway, via the apoptosis-inducing effects of caspase-3 (Yang et al., 2017). However, in hepatocytes of GIFT with knocked-down *CaSR* expression, the decreased CaSR expression levels may have prevented the accumulation of Ca^2+^ under hypoxia stress. This would explain the reduced transcript levels of *p53, TP53INP2, caspase-3*, and *caspase-8* in hepatocytes with knocked-down *CaSR* expression. Therefore, *CaSR* knockdown helped to maintain the homeostasis of the Ca^2+^ concentration inside and outside the cell, and relieve hypoxia stress-induced apoptosis.

Hypoxia-induced apoptosis is an anoxic adaptation mechanism that eliminates stressed cells (Shimizu et al., 1996). Under hypoxia stress, activation of p53 can induce cell necrosis, apoptosis, or autophagy, all of which are involved in cell death (Denko et al., 2003; Nuñez-Hernandez et al., 2018). Previous studies have shown that hypoxia stress leads to increased expression of the *p53* gene and p53 protein in oriental river prawn (*Macrobrachium nipponense*) and white shrimp (*Litopenaeus vannamei*) hepatopancreas and haemocytes, leading to apoptosis (Nuñez-Hernandez et al., 2018; Sun et al., 2016; Felix-Portillo et al., 2016). Similar results were found in this study. The transcript level of *p53* in each experimental group gradually increased under hypoxia stress. Activated endogenous expression of miR-92a in GIFT by injection of a miRNA agomir inhibited *CaSR* mRNA expression levels, thereby regulating the apoptosis response. Therefore, decreased *CaSR* expression in the agomir group may have helped to reduce the increase in p53 expression and alleviate apoptosis induced by hypoxia stress. As a key regulatory gene in the p53 regulatory network, *TP53INP2* plays an important role in the apoptosis and invasion of tumour cells. In previous studies, up-regulation of this gene was shown to induce cell apoptosis of glioma cells, breast cancer cells, and osteosarcoma cells (Yu et al., 2018; Wang et al., 2016). Up-regulation of p53 may result in a significant up-regulation of *TP53INP2*, leading to increased hepatocyte apoptosis and liver damage. However, in our study, *p53* was significantly down-regulated in the liver of GIFT during prolonged exposure to hypoxia stress. Liu et al (2005) suggested that p53 is a universal receptor for environmental stress. Therefore, down-regulation of *p53* and *TP53INP2* may be a self-adaptive mechanism under hypoxia stress in aquatic animals. After 48 h of hypoxia stress, white shrimp with silenced *p53* showed enhanced expression of cyclin-dependent kinase 2 (which is involved in cell cycle progression) and decreased caspase-3 expression, thereby regulating hepatocyte apoptosis and the cell cycle (Shimizu et al., 1996).

In our study, the transcript levels of *caspase-3* and *caspase-8* increased significantly in the liver tissues of GIFT under hypoxia stress, indicating that the caspase signalling pathway may play a role in stress adaptation. For example, apoptosis of purse red common carp (*Cyprinus carpio*) hepatocytes was found to be induced by cadmium stress, and caspase-3A activity was significantly increased in liver tissues (Gao et al., 2013). However, copper stress did not lead to up-regulation of *caspase-3* in Nile tilapia, nor did it increase the proportion of apoptotic cells, suggesting that the caspase-3-dependent or caspase-independent apoptotic pathways may not exist in this fish (Monteiro et al., 2009). In our study, up-regulated miR-92a in the agomir group inhibited the expression of the target gene *CaSR* and reduced the increase in *caspase-3* and *caspase-8* transcript levels under hypoxia stress. This may have prevented the apoptosis of GIFT liver cells under stress. In another study, down regulation of miR-532-5p expression in H9c2 cells exposed to hypoxia led to up-regulation of its target gene encoding programmed cell death protein 4, and increased the expression of caspase-3, thereby promoting hypoxia-induced apoptosis of H9c2 cells (Ma et al., 2018). Inhibition of miR-9 up-regulated the gene encoding Yes-associated protein 1, promoted cell proliferation, and inhibited apoptosis and caspase-3/7 activity in hypoxic H9c2 cells (Zheng et al., 2019).

Under normal conditions, ALT and AST in fish are mainly located in the liver, and only small amounts are released into the blood. Thus, serum ALT and AST activities are important indicators of normal liver function (Qiang et al., 2017). In other studies, increased serum ALT and AST activities in GIFT have been detected under high temperature (Bao et al., 2018) and crowding (Qiang et al., 2015). In our study, the GIFT liver may have been damaged by hypoxia stress. An increase in the membrane permeability of liver cells would lead to the release of ALT and AST, resulting in increased serum ALT and AST activities. Inhibition of *CaSR* in the agomir group may have helped to alleviate hepatocyte damage, thereby reducing serum ALT and AST activities.

In summary, up-regulation of miR-92a led to down-regulation of its target gene *CaSR* in GIFT. Stimulation of miR-92a interfered with hypoxia-induced apoptosis in hepatocytes of GIFT by targeting *CaSR*, and alleviated liver damage. Our study provides novel insights into the adaptation mechanism of GIFT to hypoxia, and suggests that miR-92a might be a target for relieving stress in farmed fish.

## Abbreviations

UTR: untranslated region
GIFT: genetically improved farmed tilapia
TP53INP2: p53-inducible nuclear protein 2
CaSR: calcium-sensing receptor
IL-1β: interleukin-1β
IL-6: interleukin-6
NC: negative control
DO: dissolved oxygen
SDS: sodium dodecyl sulphate
TBST: Tris-buffered saline with Tween
ALT: alanine aminotransferase
AST: aspartate aminotransferase
TP53INP2: tumour protein p53-inducible nuclear protein 2
GAPDH: glyceraldehyde-3-phosphate dehydrogenase

## Author Contributions

P.X. and J.H. conceived and designed the experiments; Y.F.T., J.W. B, and J.H. sampled the liver and serum tissues and extracted RNA; J.Q. constructed and analysed the miRNA; J.H.Z and T.Y.F. quantified gene mRNAs and proteins; J.W.B. analysed biochemical indicators in serum; J.Q. wrote the paper with contributions from Y.F.T., J.W.B., P.X., J.H.Z, and J.H.. All authors read and approved the final version of the manuscript.

## Acknowledgements

We thank Jennifer Smith, PhD, from Liwen Bianji, Edanz Group China (http://www.liwenbianji.cn/ac), for editing the English text of a draft of this manuscript.

## Data availability statement

All data generated or analysed during this study are included in this published article (and its Supplementary Information files).

## Additional files: supplementary figure captions

**Additional file 1: Figure S1** Original image of CaSR marker and protein level of experiment group by western blot (XLSX 1.0 MB).

